# The orientation of choroidal macrophage polarization significantly influences the development of myopia in murine models

**DOI:** 10.1101/2023.06.12.544445

**Authors:** Jing Hou, Shin-ichi Ikeda, Kiwako Mori, Heonuk Jeong, Hidemasa Torii, Kazuno Negishi, Kazuo Tsubota, Toshihide Kurihara

**Affiliations:** Department of Ophthalmology, Keio University School of Medicine, 35 Shinanomachi, Shinjuku-ku, 160-8582, Tokyo, Japan; Laboratory of Photobiology, Keio University School of Medicine, 35 Shinanomachi, Shinjuku-ku, 160-8582, Tokyo, Japan; Tsubota Laboratory, Inc., 304 Toshin Shinanomachi-ekimae Bldg., 34 Shinanomachi Shinjuku-ku, 160-0016, Tokyo, Japan

**Keywords:** Macrophage, Myopia, Choroid, Vascular Maintenance, Anti-inflammatory.

## Abstract

Myopia is a primary contributor to visual impairment and has emerged as a global public health concern. Evidence indicates that one of the main structural features of myopia is the corresponding decrease in choroidal thickness, and choroidal macrophages play an important role in maintaining the choroidal thickness. Nevertheless, the effect of choroidal macrophages on myopia remains unclear. Here, we discovered that the continuous intraperitoneal injection of clodronate liposomes depleted choroidal macrophages and leads to myopia, which confirmed that the presence of choroidal macrophages plays an important role in myopia development. Subsequently, based on the phenotypic characteristics of macrophages, experiments were designed to study the effects of different polarization directions of macrophages on myopia development. We found that lipopolysaccharides (LPS) injection can induce the polarization of choroidal M1 macrophages, thinning the choroidal thickness and resulting in myopia. Conversely, IL-4 or IL-13 injection causes choroidal M2 macrophage polarization, thickens the choroid, and suppresses the progression of myopia. Additionally, we demonstrated that the opposite effects of M1 and M2 macrophages on myopia development may be related to their impacts on choroidal thickness, inflammation, and oxidative stress response. These findings establish that choroidal macrophages are critically important in the development of myopia and provide new strategies for the development of myopic therapies.

## Main Text

### Introduction

Myopia, commonly referred to as “short-sightedness,” is the most prevalent vision-related disease, which is caused by refractive error depending on the elongation of the eyeball along the visual axis (1–4). Current research has shown that myopia can be influenced by a variety of factors, including light environmental impacts, dietary factors, and genetics; however, the pathogenesis of myopia remains unclear (5–8). Numerous studies have suggested that inflammation plays a role in the development of myopia: increased inflammation in the eye accelerates the development of myopia, whereas lower inflammation inhibits it (9–12). It is worth noting that macrophages are essential for the occurrence, maintenance, and resolution of inflammation (13).

Macrophages are an essential component of the innate immune system that exists in all mammalian tissues and exhibit great functional diversity. It not only plays a variety of functions related to inflammation but also plays a crucial role in the development and maintenance of homeostasis (14, 15). A healthy cornea contains a population of macrophages that maintain corneal lymphatic homeostasis and promote lymphangiogenesis under inflammatory conditions (16). The neural retina is populated by microglia, specialized resident macrophages that contribute to the structural and functional integrity of neuronal synapses (16, 17). Under homeostatic conditions, there are no microglia in the subretinal space or the choroid and are restricted to the inner layers of the retina (18–20). Further, the choroid was found to contain a dense network of resident tissue macrophages as early as the 1990s (21). At the level of choroidal arteries and arterioles, dendritic resident macrophages were closely aligned along the vessel wall. In the choroidal terminal choriocapillaris, they are not tightly packed on the vessel wall, but are widely distributed in the connective tissue between blood vessels (22). These findings provide a basis for exploring the relationship between choroidal vessels and macrophages. Subsequently, researchers found that the ablation of a considerable majority of choroidal macrophages was associated with broad and gradual atrophy of the choroidal vasculature and thinning of the choroid (23, 24). This finding reveals that choroidal macrophages play a hitherto underestimated role in the maintenance of the choroidal vascular structure. In the field of myopia research, the stability of the choroidal vasculature is recognized as the most important component for maintaining choroidal thickness, and choroidal thickness variation is a structural characteristic of the development of myopia (25–28). Therefore, we hypothesized that choroidal macrophages play a crucial role in the progression of myopia.

Macrophage subsets can be classified as classic M1 macrophages and alternative M2 macrophages, according to their pro- or anti-inflammatory properties in the polarized state (29). M1 macrophages can be polarized by lipopolysaccharide (LPS), leading to a high expression of inflammatory cytokines, such as tumor necrosis factor-α (TNFA) and interleukin-6 (IL6), as well as reactive oxygen species. Correspondingly, M2 macrophage polarization induced by IL-4 and IL-13 diminishes and alleviates inflammation following infection or damage, and inhibits the production of reactive oxygen species (30–32). Studies have shown that inflammatory diseases increase the risk of myopia, and inflammation in the eye directly contributes to the progression of myopia (12, 33), all this underscores the importance of controlling inflammation in inhibiting the development of myopia. In addition, it has been suggested that the existence of several supplements which convert M1 into M2 macrophages, but this property has not been studied and discussed in conjunction with myopia suppression (34–36). Therefore, we hypothesized that different subsets of macrophages in the choroid have diametrically opposite effects on the development of myopia according to their polarization direction and characteristics.

In this study, we investigated the depletion and polarization of choroidal macrophages in a mouse model and combined a lens-induced myopia model (LIM) with the polarization of choroidal macrophage to investigate the effect of different macrophage polarization directions on myopia development. We found that choroidal macrophage ablation was associated with myopia progression. We also discovered that the tendency for myopia development or inhibition is closely related to the direction of macrophage polarization in the choroid and that macrophages interfere with the development of myopia based on their effects on inflammation and oxidative stress.

Taken together, our findings suggest that macrophages play a key role in myopia and emphasize the different characteristics of macrophages as a potential mechanism for developing or inhibiting myopia. Regulating the phenotype of macrophages in the choroid may have therapeutic effects on maintaining choroidal vasculature thereby slowing the progression of myopia.

## Results

### Sustained intraperitoneal injection of clodronate liposome depletes macrophages in the choroid and can trigger myopia

It has been previously reported that resident macrophages in the choroid positively affect the choroidal thickness and maintain vascular status (24). In contrast, a series of animal models have demonstrated that choroidal thinning is an inevitable change in the development of experimental myopia and is a prominent feature in the development of myopia (26, 28, 37–39). To investigate the effect of choroidal-resident macrophages on myopia in a mouse model, 3-weeks-old C57BL/6 J mice were randomly divided into 2 groups and were injected intraperitoneally with clodronate or phosphate-buffered saline (PBS) liposomes. In the clodronate liposome group, after being engulfed by macrophages, the clodronate liposomes were degraded by lysosomal phospholipases, which released clodronate into the macrophages and induced apoptosis, thereby promoting macrophage depletion (40). The injection volume of clodronate liposome was 0.1 ml per 10 g of mice and the injection frequency was once every two days. Refraction, axial length, and choroidal thickness were measured using an infrared photorefractor and a spectral domain-optical coherence tomography (SD-OCT) system, before injection and after 8 days of injections, respectively (Fig. 1A). Mice injected with clodronate liposome showed a higher refractive shift (−7.30 ± 3.61 D vs. +1.17 ± 4.40 D, *P* < 0.001), axial length elongation (0.08 ± 0.02 mm vs. 0.05 ± 0.02 mm, *P* < 0.01), and thinner choroid (−1.56 ± 0.89 mm vs. 1.09 ± 0.51 mm, *P* < 0.001) compared to the PBS liposome injection group (Fig. 1B). Flow cytometry analysis of the percentage of choroidal resident macrophages demonstrated a corresponding decrease after 8 days of clodronate liposome injection compared to the control group (2.583 ± 0.002% vs. 9.290 ± 0.018%, *P* < 0.05) (Fig. 1C). These results indicated that clodronate liposome injection depletes resident macrophages in the choroid, promotes choroidal thinning, and induces myopia in 3-week-old mice. Previous studies showed that choroidal thickness did not increase and tended to be stable from approximately 8 weeks of age in mice (41). However, whether the depletion of choroidal macrophages in adult mice also contributes to myopia has not been elucidated. To rule out the possibility that choroidal macrophage depletion can cause myopia independent from the developmental stage of choroidal thickness, 8-weeks-old adult mice were injected with clodronate liposomes or PBS liposomes in the same manner, and changes in refraction, axial length, and choroidal thickness were analyzed 8 days later (Fig. 1D). Intraperitoneal injection of clodronate liposomes significantly reduced body weight while consuming resident macrophages, which affects the comparison of changes in the axial length of the eye. To avoid the effects of body weight changes on the axial length, we used the ratio of the axial length to body weight to represent the changes in the axial length of the eye. In the clodronate liposome injection group, a higher refraction shift (−7.52 ± 2.57 D vs. +1.52 ± 4.91 D, *P* < 0.01), a higher axial length/body weight (0.168 ± 0.015 mm/g vs. 0.155 ± 0.002 mm/g, *P* < 0.05), and thinner choroidal (−2.39 ± 1.33 mm vs. −0.12 ± 1.48 mm, *P* < 0.01) were observed compared to the PBS liposome injection group (Fig. 1E), suggesting that clodronate liposome injection causes choroidal resident macrophage depletion and thinning of the choroid, which leads to myopia regardless of whether the choroid is developing.

**Figure 1.**
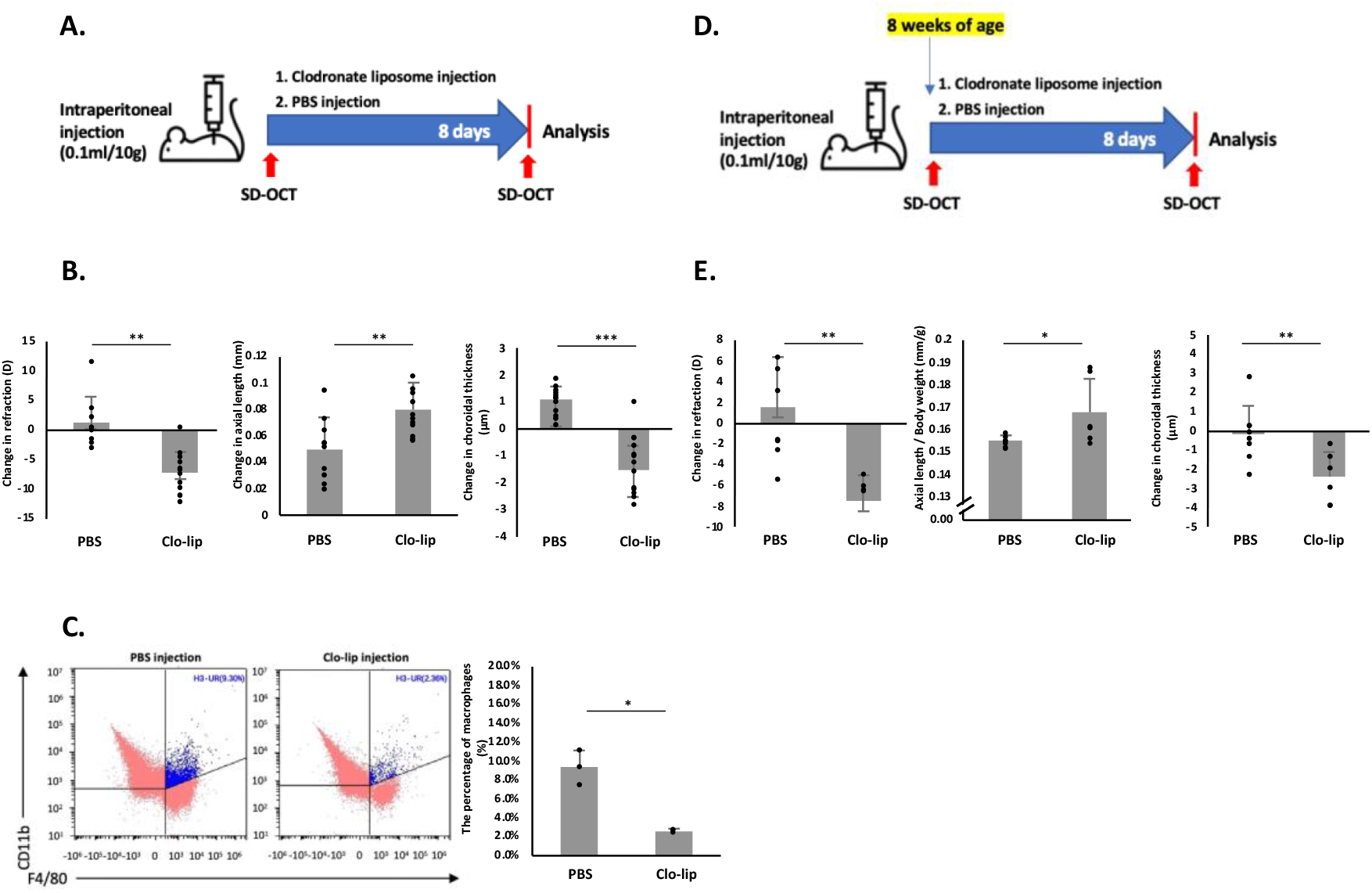
Myopia is accompanied by sustained depletion of choroidal-resident macrophages in mice at 3 and 8 weeks of age. (**A**) Three-weeks-old wild-type C57BL/6JJc1 mice were injected with either 0.1 ml/10 g body weight clodronate liposome or control liposome once every two days (n=12 per group). Refraction, axial length, and choroidal thickness were measured at baseline and 8 days after 4 injections with an infrared photorefractor and SD-OCT system. **(B)** Compared to the control group, the clodronate liposome (clo-lip) injection group showed a larger refractive change (*P* < 0.001), axial eye growth (*P* < 0,01), and thinner choroid (*P* < 0.001). Bars represent mean +/− standard deviations. **(C)** The percentage of macrophages in the choroid was analyzed using flow cytometry. Macrophages were identified as CD11b^+^ and F4/80^+^, appearing in the H3-UR region. Compared with the control liposome injection group, the proportion of macrophages in the clodronate liposome injection group was significantly reduced (*P* < 0.05). **(D)** Eight-weeks-old wild-type C57BL/6JJc1 mice (4 per group unless otherwise indicated) were injected with 0.1 ml/10 g body weight clodronate liposome and control liposome once every two days. Refraction, axial length, and choroidal thickness were measured at baseline and 8 days after 4 injections with an infrared photorefractor and SD-OCT system. **(E)** Compared to the control group, the clodronate liposome (clo-lip) injection group showed a remarkably larger refractive change (*P* < 0.01), axial length elongation (*P* < 0.05), and a thinner choroid (*P* < 0.01). Bars represent mean +/− standard deviations. Differences between the groups were compared using a t-test. \**P* < 0.05. \*\**P* < 0.01. \*\*\**P* < 0.001.

### Continuous intraperitoneal injection of LPS stimulates choroidal M1 macrophage polarization and induces myopia in a murine model

Macrophages are classified into two categories based on their activation state and functions: M1 type (classically activated macrophages) and M2 type (alternatively activated macrophages), which exhibit pro-inflammatory and anti-inflammatory properties, respectively (31). As the upregulation of allergic inflammation has been implicated in the progression of myopia in animal models (33), we investigated whether M1 macrophage polarization can induce myopia in a mouse model. Mice were divided into LPS (10 mg/ml) and PBS injection groups for daily intraperitoneal injections. To determine whether myopia had developed, refraction, axial length, and choroidal thickness were measured using SD-OCT before the injections, 1 week after, and 2 weeks after the injections (Fig. 2A). After 1 week of LPS injection, LPS-injected mice showed refractive shift (−9.21 ± 5.91 D vs. +2.91 ± 3.91 D, *P* < 0.001), rise in axial length/body weight (0.28 ± 0.01 mm/g vs. 0.22 ± 0.005 mm/g, *P* < 0.001), and thinning of the choroid (−1.11 ± 0.90 mm vs. 1.36 ± 0.70 mm, *P* < 0.001), compared to mice injected with PBS for a week. The injections were continued until the end of the 2 weeks, and LPS-injected mice maintained a significantly higher refractive shift (−7.23 ± 4.36 D vs. +8.41 ± 7.01 D, *P* < 0.001), increase in the ratio of axial length to mouse body weight (0.20 ± 0.005 mm/g vs. 0.17 ± 0.004 mm/g, *P* < 0.01), and thinning of the choroid (−1.77 ± 1.02 mm vs. 2.05 ± 0.66 mm, *P* < 0.001), in comparison to PBS-injected mice (Fig. 2B). Choroidal blood perfusion was measured by optical coherence tomography angiography (OCTA), and the results showed that the choroidal blood perfusion of mice injected with LPS for 2 weeks was reduced compared to that of mice injected with PBS (37.36 ± 3.74 % area vs. 46.79 ± 4.74 % area, *P* < 0.001) (Fig. 2C). These data show that the intraperitoneal injection of LPS can induce myopia in a mouse model. Choroid samples were collected for real-time polymerase chain reaction (PCR) analysis to verify the polarization of M1 macrophages in the choroid stimulated by intraperitoneal injection of LPS. The expression levels of the pro-inflammatory cytokines *Tnf* and *Il6* were significantly upregulated in LPS-injected choroid samples compared to those in PBS-injected choroid samples (Fig. 2D). These data indicate that the intraperitoneal injection of LPS polarizes M1 macrophages in the choroid, promotes the release of inflammatory cytokines, and causes myopia in a mouse model. As reported in LPS-treated macrophages, pro-inflammatory cytokines (TNFA, and IL6) secretion was significantly attenuated by eicosapentaenoic acid (EPA), which can inhibit myopia progression (42, 43). Therefore, an experiment was designed to administer an EPA diet to LPS-induced myopic mice, and it was found that myopia did not progress (Supplementary Figs. 1A, B, C). Taken together, these data suggest that the LPS-induced phenotypic switch toward M1 macrophages is associated with the development of myopia.

**Figure 2.**
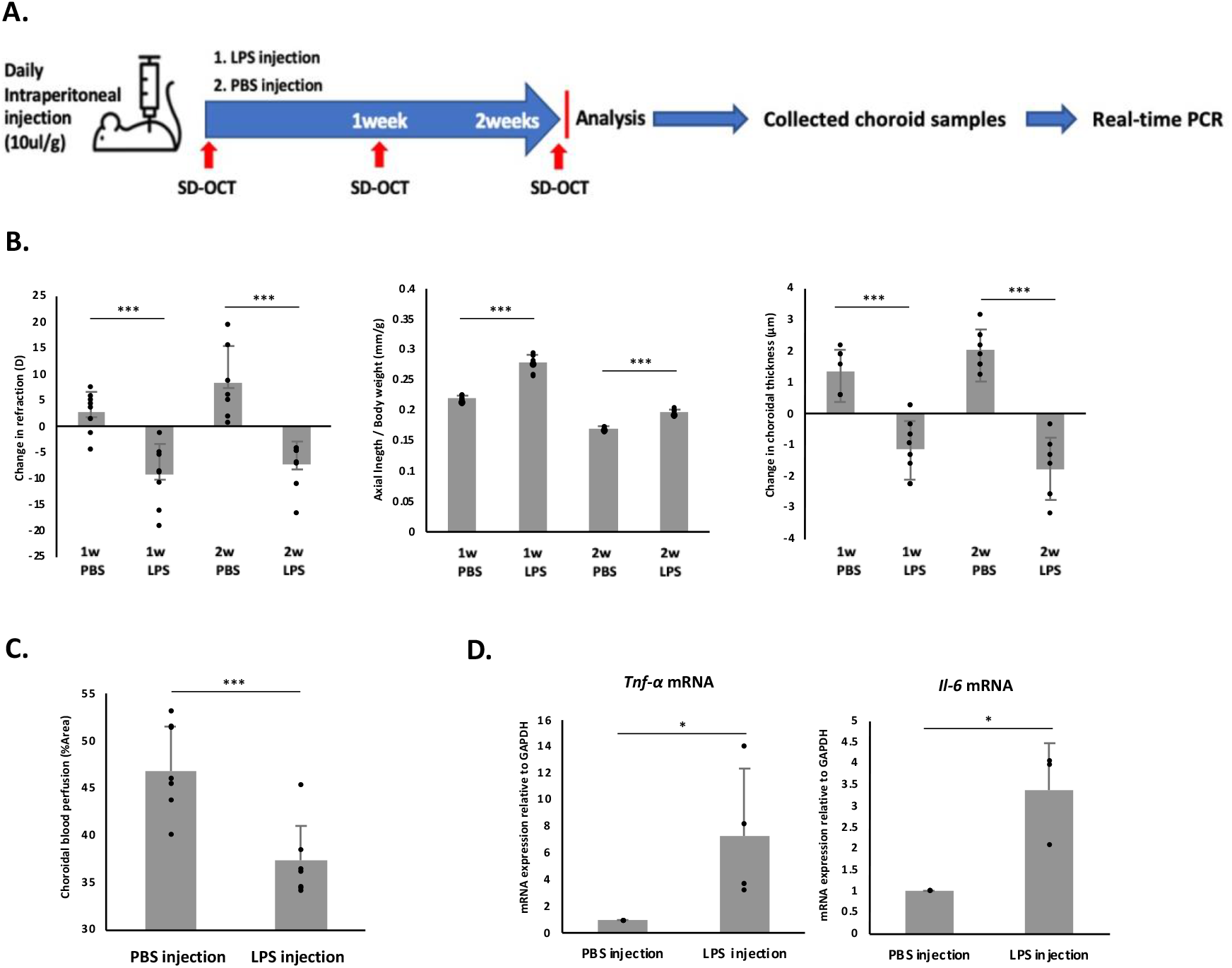
The polarization of M1 macrophages in the choroid can induce myopia in a murine model. **(A)** Three-weeks-old wild-type C57BL/6JJc1 mice were divided into two groups (4 per group unless otherwise indicated) and one group was intraperitoneally injected with 10 ul/g LPS and the other with PBS. The injection lasted for 2 weeks, and refraction, axial length, and choroidal thickness were measured weekly with an infrared photorefractor and SD-OCT system. Choroidal blood perfusion was measured with OCTA. In addition, choroid samples were collected for real-time PCR. **(B)** Compared to the PBS injection group, the LPS injection group showed a significantly larger refractive change (*P* < 0.001), significant axial eye growth (*P* < 0,001), and substantially thinner choroid (*P* < 0.001). Bars represent mean +/− standard deviations. **(C)** The choroidal blood perfusion decreased significantly after 2 weeks of LPS injection, compared to the PBS injection group. **(D)** mRNA levels of *Tnf-α* and *IL-6* significantly increased after 2 weeks of LPS injection. *P* values were obtained from a t-test. \**P* < 0.05. \*\**P* < 0.01. \*\*\**P* < 0.001.

### The constant intraperitoneal injection of IL-4/IL-13 encouraged the polarization of choroidal M2 macrophages and suppressed myopia progression in a murine model of LIM

We found that intraperitoneal injection of LPS enhanced the expression of pro-inflammatory cytokines in the choroid and promoted myopia development; contrarily, choroidal M2 macrophage polarization should inhibit myopia development. Previous studies have demonstrated that IL-4 and IL-13, as anti-inflammatory interleukins, can directly activate M2 macrophages, which can be identified by the expression of M2 macrophage markers, including mannose receptor 1 (Mrc1), also referred to as CD206 and CD163 (44). Above all, to investigate the frequency of continuous IL-4 injection, we compared the proportion of M2 macrophages in the choroid at 0 h, 24 h, and 48 h after being subjected to 0.1 µg/100 µl IL-4 intraperitoneal injection; F4/80 and CD11b-positive cells were regarded as macrophages and CD206 was used as a marker to identify M2 macrophages. Flow cytometry analysis showed a growth in the ratio of macrophages (6.73 ± 0.22 % vs. 3.78 ± 0.41 %, *P* < 0.001) and M2 macrophages (43.57 ± 4.72 % vs. 30.33 ± 8.35 %, *P* < 0.05) at 48 h after IL-4 intraperitoneal injection, compared to 0 h (Fig. 3A). Real-time PCR detection after IL-4 injection under the same conditions showed that *Mrc1* and *CD163* mRNA expression in the choroid was enhanced 48 h after IL-4 injection (Fig. 3B). These results showed that the optimal frequency of IL-4 injection is once every two days. Next, we examined whether the promotion of M2 macrophage polarization by IL-4 injection on alternate days could inhibit myopia progression in a LIM mouse model. All 3-weeks-old mice were subjected to LIM, in the same manner as previously established (45), and administered with either PBS or 0.1 ug/100 ul of IL-4 injection until 6 weeks of age (Fig. 3C). In the PBS injection group, eyes treated with −30 diopter (D) lenses showed refractive change (−11.52 ± 4.91 D vs. +12.45 ± 8.19 D, *P* < 0.01), increased axial length (0.22 ± 0.01 mm vs. 0.18 ± 0.01 mm/g, *P* < 0.05), and decreased choroidal thickness (−1.74 ± 0.66 mm vs. 3.91 ± 0.95 mm, *P* < 0.001) compared to eyes treated with 0D lenses. In contrast, the IL-4 injection group showed smaller refractive change (−1.39 ± 2.91 D vs. −11.52 ± 4.91 D, *P* < 0.05), smaller axial elongation (0.19 ± 0.01 mm vs. 0.22 ± 0.01 mm/g, *P* < 0.05), and thickening of the choroid (1.39 ± 0.19 mm vs. −1.74 ± 0.66 mm, *P* < 0.001) in eyes with -30D lenses, compared to the PBS-injected group with −30D lenses (Fig. 3D). Meanwhile, to demonstrate the influence of continuous IL-4 injection on choroidal blood perfusion, 3-weeks-old mice were divided into three groups: control 0D group (both eyes were treated with 0D lenses), control −30D group (binocular myopia induction), and IL-4 −30D group (binocular myopia induction with IL-4 injection once every two days). OCTA was used to measure choroidal blood perfusion at the beginning (3 weeks old) and end (6 weeks old) of myopic induction. The results showed that choroidal blood perfusion was lower (8.59 ± 5.90 %area vs. −6.29 ± 8.73 %area, *P* < 0.01) after 3 weeks of myopia induction, compared to the control 0D group. Nevertheless, in the IL4 −30D group, choroidal blood perfusion was improved (−6.29 ± 8.73 %area vs. 1.54 ± 9.34 %area, *P* < 0.05), compared to the control -30D group (Fig. 3E). To visualize the polarization of M2 macrophages, after measuring choroidal blood perfusion at the end stage, choroid samples were collected for flow cytometry, and data analysis revealed the corresponding growth in the proportion of M2 macrophages (46.52 ± 0.05% vs. 35.71 ± 0.06%, *P* < 0.05) in the IL-4 injection group with −30D lenses, compared to the PBS injection group with −30D lenses (Fig. 3F). Collectively, these findings demonstrate that IL-4 injection can stimulate the polarization of M2 macrophages in the choroid and suppress the development of myopia by improving choroidal blood perfusion in a LIM mouse model. To reinforce the conclusion that M2 macrophages can inhibit myopia progression, we performed the same experiments with 0.1 µg /100 µl IL13 injection instead of IL-4 injection, and the results also showed that M2 macrophages were polarized 48 h after IL-13 injection (Supplementary Figs. 2A, B). The results of the LIM mouse model combined with IL-13 injection were as expected; IL-13 also polarized M2 macrophages, suppressed myopia progression, and improved choroidal blood perfusion in a murine model of LIM (Supplementary Figs. 2C, D, E, F).

**Figure 3.**
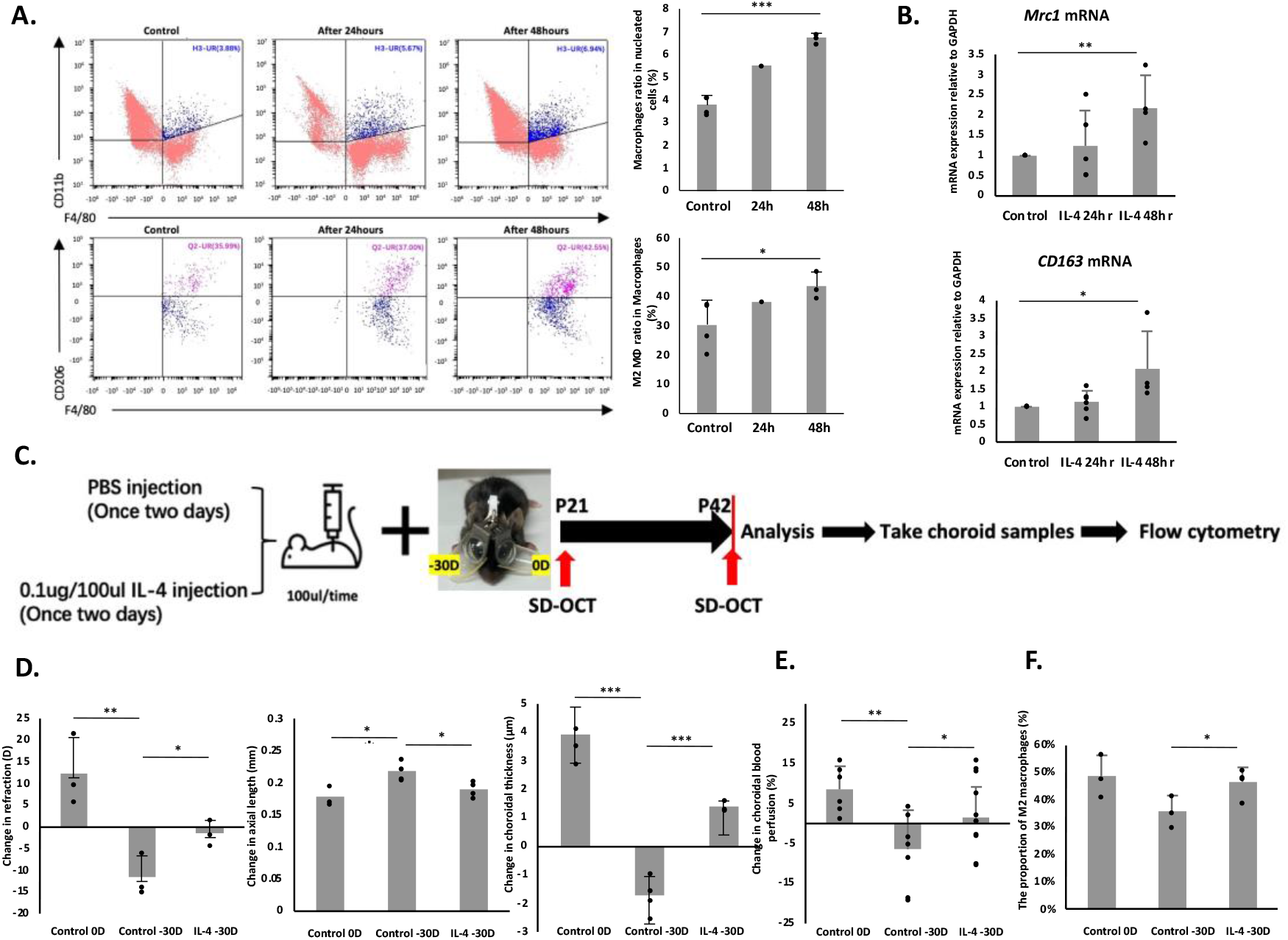
Inhibition of myopia progression by choroidal M2 macrophage polarization was demonstrated in a lens-induced myopia mouse model. Three-week-old wild-type C57BL/6JJc1 mice were injected with 0.1 ug/100 ul IL-4, and choroid samples were collected at 0 h, 24 h, and 48 h after injection for flow cytometry and real-time PCR. **(A)** Macrophages in the H3-UR region were denoted as CD11b^+^ and F4/80^+^. The proportion of macrophages increased significantly after IL-4 injection (*P* < 0.001). M2 macrophages in the Q2-UR were identified as CD206^+^. At 48 h after IL-4 injection, M2 macrophages were dramatically polarized (*P* < 0.05). **(B)** Real-time PCR results showed that the mRNA expression levels of *Mrc1* and *CD163* increased significantly 48 h after IL-4 injection. **(C)** Three-week-old wild-type C57BL/6JJc1 mice were divided into two groups and injected with either 0.1 ug/100 ul IL-4 or PBS on alternate days with myopia induction (4 per group unless otherwise indicated). Refraction, axial length, and choroidal thickness were measured at the initial (3-week-old) and final (6-week-old) stages of myopic induction using an infrared photorefractor and SD-OCT system. **(D)** Eyes treated with −30D lenses showed a significantly larger refractive change (*P* < 0.01), greater axial length elongation (*P* < 0.05), and increased choroidal thickness (*P* < 0.001) than those treated with 0D lenses in the PBS injection mice. Mice injected with 0.1 ug/100 ul IL-4 showed a significantly smaller refractive change (*P* < 0.05), smaller change in axial length(*P* < 0.05), and positive change in choroidal thickness with a statistical significance (*P* < 0.001). Bars represent the mean +/− standard deviation. **(E)** To evaluate changes in choroidal blood perfusion, three-weeks-old mice were divided into three groups: both eyes with 0D lenses (control 0D group), binocular myopia induction (control -30D group), and binocular myopia induction with IL-4 injection once every two days group (IL-4 -30D group). Choroidal blood perfusion was assessed at the initial (3-week-old) and end (6-week-old) stages of myopic induction using OCTA. Compared to the control 0D group, choroidal blood perfusion was significantly decreased in the control -30D group. Accordingly, compared with the control -30D group, choroidal blood perfusion was remarkably improved in the IL-4 -30D group. **(F)** Mice used to measure choroidal blood perfusion were euthanized at 6 weeks of age, and the choroids were collected for flow cytometry. The results showed that mice treated with -30D lenses and injected with IL-4 had a significant proportion of M2 macrophages compared to those treated with -30D lenses and injected with PBS. *P* values correspond to comparisons made using one-way ANOVA. \**P* < 0.05. \*\**P* < 0.01. \*\*\**P* < 0.001.

### Berberine, an eye drop that polarizes M2 macrophages, inhibits myopia progression after consistent use

After realizing that M2 macrophage polarization may inhibit the development of myopia, we used an M2 macrophage-activating drug as an eye drop to verify its effect on the development of myopia. Some studies have shown that berberine promotes the polarization of M1 macrophages to the M2 anti-inflammatory phenotype in mice (46, 47). To verify the role of berberine eye drops in the development of myopia, we examined and compared changes in refraction, axial length, and choroidal thickness in 3-weeks-old C57BL/6 J mice after myopia induction for 3 weeks and the administration of either berberine or dimethyl sulfoxide (DMSO) eye drops. The control −30D group (received −30D lenses for both eyes) and 2% DMSO −30D group (binocular myopia induction with the application of 2% DMSO eye drops daily) showed a large refractive change (*P* < 0.001), increased axial length elongation (*P* < 0.05), and decreased choroidal thickness (*P* < 0.001), compared to the control 0D group (received 0D lenses for both eyes). In addition, the berberine −30D group (binocular myopia induction with daily 0.26 mg/ml eye drops) exhibited smaller refractive changes (2.12 ± 1.63 D vs. −5.25 ± 2.89 D, *P* < 0.001), smaller axial elongation (0.19 ± 0.02 mm vs. 0.22 ± 0.04 mm/g, *P* < 0.05), and a relatively thicker choroid (0.79 ± 0.64 mm vs. −1.56 ± 0.76 mm, *P* < 0.001), compared to the 2% DMSO −30D group. These results indicate that drugs used to activate the polarization of M2 macrophages may also have the potential to inhibit the development of myopia (Supplementary Figs. 3A, B, C).

### Effects of LPS and IL-4 injections on nicotinamide adenine dinucleotide phosphate (NADPH) oxidation activity

We inferred that the direction of macrophage polarization may influence the development or inhibition of myopia through its pro-inflammatory or anti-inflammatory properties. However, the other mechanisms underlying this association remain largely unknown. LPS-treated M1 macrophages are associated with high levels of reactive oxygen and nitrogen species (ROS and RNS, respectively) and matrix metalloproteinase (MMP) activity, and ROS production is critical for M1 macrophage activation and function (48, 49). Correspondingly, ROS overproduction may impair blood perfusion to the retina, and circulatory disturbances during the development of myopia can lead to oxidative stress (50, 51). To investigate whether M1 macrophage polarization also induces myopia by promoting ROS and oxidative stress response, and whether M2 macrophage polarization suppressed myopia progression by inhibiting ROS release and oxidative stress response, the mRNA expression of NADPH oxidase 2 (*Nox2*), which is the most important source of ROS, and the expression of *Nox4, Mmp2*, and *Mmp9* in the choroid were observed after either LPS or IL-4 injection (Fig. 4A). Quantitative analysis showed increased *Nox2, Nox4, Mmp2,* and *Mmp9* mRNA expression in the LPS injection group compared to that in the PBS injection group (Fig. 4B). In contrast, 48 h after IL-4 injection, the mice showed lower *Nox2, Mmp2,* and *Mmp9* mRNA expression (Fig. 4C). Since 8-hydroxy-2-deoxyguanosine (8-OHdG) is an effective marker for assessing oxidative DNA damage (52), we investigated the level of 8-OHdG in the choroid after intraperitoneal injection of either LPS or IL-4 and found that the IL-4 injection group showed a reduction in choroidal 8-OHdG concentration compared to the control group (*P* < 0.01). In contrast, the concentration of choroidal 8-OHdG was higher in the LPS injection group, compared to the IL-4 group (*P* < 0.05) (Fig. 4D). Taken together, these data suggest that polarized M1 macrophages may induce myopia by promoting oxidative stress, whereas polarized M2 macrophages may inhibit myopia progression by suppressing the oxidative stress response.

**Figure 4:**
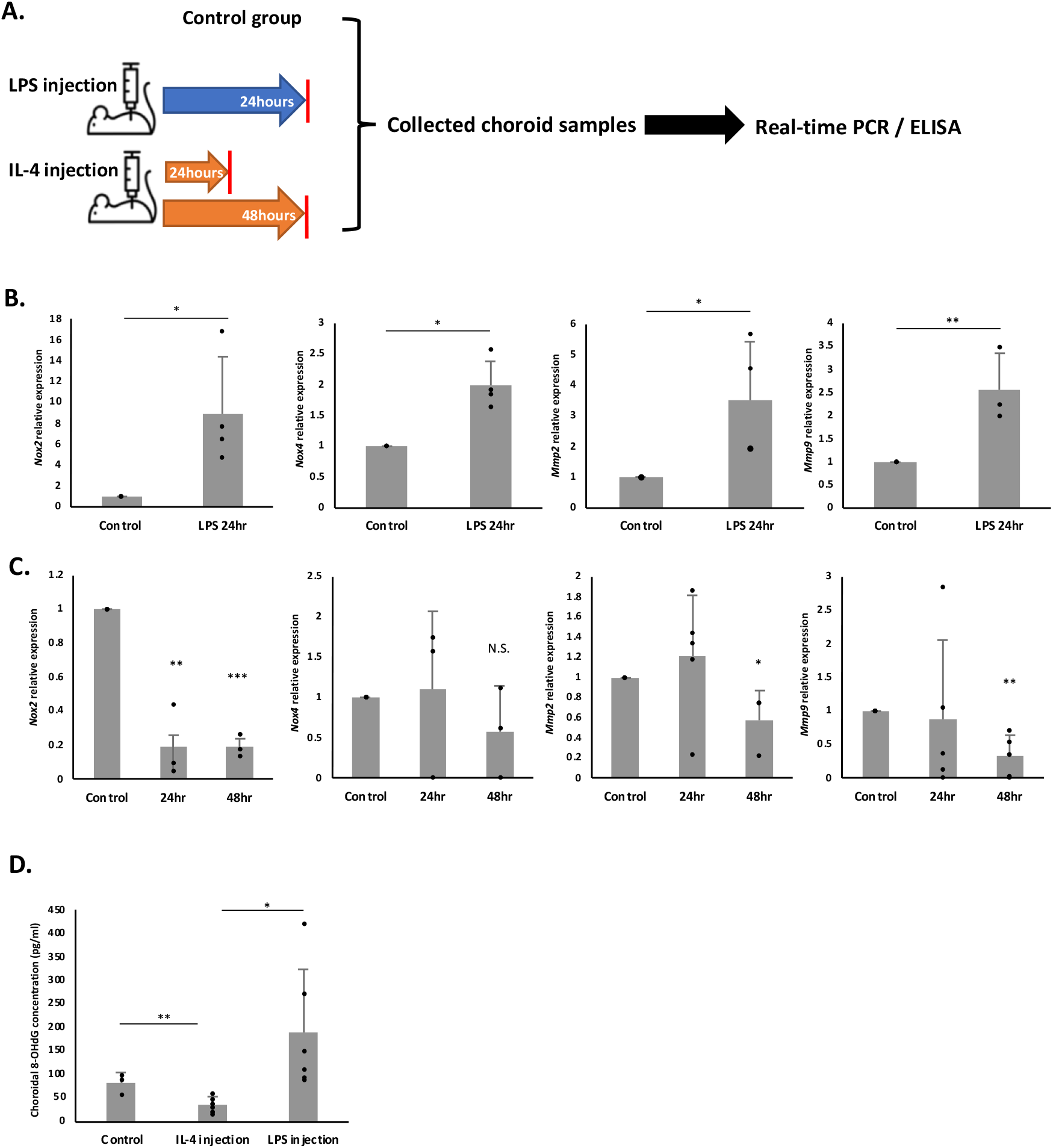
Effect of macrophage polarization direction on oxidative stress response. **(A)** Experimental plan for the assessment of *Nox2, Nox4, Mmp2,* and *Mmp9* mRNA expression in the choroid during M1 macrophage polarization and M2 macrophage polarization. Three-weeks-old wild-type C57BL/6JJc1 mice were injected with LPS to polarize M1 macrophages and with IL-4 to polarized M2 macrophages, while the control group was injected with PBS. Choroid samples were collected 24 h after LPS injection, 24 h and 48 h after IL-4 injection, and after PBS injection for real-time PCR and ELISA experiments. **(B)** Comparison of *Nox2, Nox4, Mmp2,* and *Mmp9* mRNA expression in the choroid between the control and M1 macrophage polarization group showed a significant increase in the LPS-injected group. **(C)** Quantitative PCR analyses of the choroid 48 h after IL-4 injection indicated a corresponding reduction in the expression of *Nox2, Mmp2,* and *Mmp9* mRNA, corresponding to M2 macrophage polarization. **(D)** ELISA analysis results showed that compared to the control group, the choroidal 8-OHdG concentration was decreased in the IL-4 injection group (*P* < 0.01). On the contrary, the choroidal 8-OHdG concentration was increased in the LPS injection group, compared to the IL-4 injection group (*P* < 0.05). *P* values indicate comparisons relative to the control, one-way ANOVA, or t-test. \**P* < 0.05. \*\**P* < 0.01. \*\*\**P* < 0.001.

## Discussion

In this paper, we have found that M1 macrophage accelerates myopia while M2 macrophage suppressed myopia progression, this statement is supported by the data performed and described. Since the choroid is now considered to be the most important tissue for the development of myopia and maintenance of the choroidal vasculature is critical for the suppression of myopia. Our findings on the role of macrophages are indeed a novel concept and open up new avenues for further investigation.

Macrophages are tissue-resident or infiltrating immune cells that are present in all tissues. In addition to their immune properties, macrophages play an important role in regulating normal tissue development and homeostasis (14). Researchers have found that resident macrophages are densely distributed in the choroid tissue, a connective tissue rich in pigments and blood vessels that supply blood and nutrients to the outer retina (16, 53). However, the precise role of choroidal-resident macrophages has rarely been directly studied. Recently, choroidal macrophages have been suggested to perform an underappreciated trophic function by preserving the vascular structure of the choroid (24). In the present study, we confirmed that choroidal thinning occurred following depleted choroidal-resident macrophages and was accompanied by axial length elongation and a significant refractive shift. This unexpected finding supports the previously unknown concept that macrophages in the developing choroid are associated with choroidal growth and myopia.

The ability of resident immune cells to maintain the vasculature changes depending on the developmental state of the tissue (24). Microglia, as specialized macrophages, play an important role in shaping the central nervous system (CNS) vasculature during development but are not necessary for the maintenance of the mature uninjured CNS (17, 54). The choroidal thickness steadily increases and reaches a steady state at 7–9 weeks of age; thereafter, it gradually decreases with age (41). In our experiments, we used 8-weeks-old mice with a plateau in choroidal thickness, depleted choroidal-resident macrophages, and found that the choroid became thinner and myopia still occurred. These results demonstrate that choroidal macrophages effectively support choroidal growth during development and maintain choroidal stability after maturation.

Studies of the normal choroid in mice have revealed a network of F4/80^+^ (CD160) and CD169^+^ macrophages widely distributed around choroidal blood vessels (55). F4/80 is one of the most specific cell surface markers of murine macrophages, and CD169^+^ macrophages are a unique subset of macrophages involved in multiple immune responses (56, 57). Activated macrophages can be broadly divided into the M1 and M2 cell subsets, representing the two extremes of the macrophage activation spectrum that can occur *in vivo* (58). M1 macrophage polarization occurs in many ocular inflammatory diseases and is accompanied by the production of pro-inflammatory cytokines; these inflammatory reactions induce local biochemical reactions, leading to local tissue remodeling and disease progression. Numerous studies have shown that inflammatory diseases of the eye can increase the risk of myopia. Myopic refractive errors are significantly increased in patients with juvenile chronic arthritis (JCA)-related uveitis, and inflammatory cytokines IL6 and TNFA were found to be overexpressed in the aqueous humor of uveitis patients (9, 59). Multifocal choroiditis (MFC) was definitely associated with myopia, and statistical analysis of 41 MFC patients showed that their average refractive error was -2.19 diopters (60). These findings suggest that acute or chronic systemic inflammatory diseases increase the risk of myopia. In this study, we demonstrated that LPS injection stimulates the expression of pro-inflammatory cytokines in the choroid, reduces choroidal thickness, and induces myopia in a murine model. This study suggests that the pro-inflammatory properties of M1 macrophages may contribute to the pathogenesis of myopia.

In contrast, some anti-inflammatory substances have been shown to inhibit the development of myopia when administered orally or as eye drops. Atropine eye drops are currently the most effective therapy for myopia control and have been shown to be associated with effective downregulation of eye inflammation (12, 61). Resveratrol eye drops reduced inflammation in a form-deprivation myopia (FDM) animal model to prevent myopia development (62). In oral preparations, the oral administration of lactoferrin (LF), which has anti-inflammatory properties, inhibited myopia progression in a LIM mouse model (63). However, no studies have shown how the polarization of M2 macrophages with anti-inflammatory properties directly affects the development of myopia. Based on the highly plastic nature of macrophages, we administered IL-4 and IL-13 injections to induce the polarization of M2 macrophages in the choroid and found that the expression of anti-inflammatory cytokines was activated in the choroid and inhibited the progression of myopia. Taken together, these results indicate that direct stimulation of M1 macrophages to release inflammatory factors induces inflammatory reactions, leading to accelerated myopia, which can be reversed by direct stimulation of M2 macrophages to release anti-inflammatory factors.

We also found that with the activation of M2 macrophages, choroid blood perfusion, and choroid thickness also increased correspondingly, this indicates M2 macrophages in the choroid play a crucial role in choroidal thickness and maintenance of blood perfusion. This is consistent with its property of maintaining the stability of the vascular structure. Under a physiological state, M2 macrophages contribute to endothelial barrier protection and facilitate vascular stability and maturation (64, 65). In a pathological condition, ameliorating macrophage inflammation or polarizing it into the M2 phenotype can prevent the rupture of intracranial aneurysms (66). M2 macrophage polarization in the aneurysmal aortic wall heals the aortic wall and maintains vascular integrity as the disease progresses (67). The ability of M2 macrophages to maintain vascular stability provides strong evidence that it can maintain the stability and overall morphology of the choroid vascular structure and thus against the development of myopia.

In addition to influencing myopia development through their anti-inflammatory or maintaining vascular structural stability properties, M2 macrophages may also inhibit the progression of myopia through their ability to inhibit oxidative stress. In contrast, the mechanism of myopia induced by M1 macrophage polarization may be related to the activation of oxidative metabolism. Polarized M1 macrophages produce reactive oxygen species (ROS) and drive inflammatory responses, leading to the production of high levels of nitric oxide (NO) and NO synthase (iNOS); excessive ROS can cause oxidative stress and damage biomolecules. Oxidative stress upregulates the expression of matrix metalloproteinases-2 and -9 (MMP-2 and MMP-9) (68–71). M2 macrophages have the opposite effects (72). Our study demonstrated that stimulating the polarization of M1 macrophages in the choroid induces the expression of NADPH oxidases, matrix metalloproteinase, and 8-OHdG while stimulating the polarization of M2 macrophages results in the exact opposite effect. Moreover, there is evidence that oxidative stress plays a role in the development of myopia-associated disorders at the cellular level (51). In pathological myopia, retinal hypoxia and ischemia lead to the accumulation of ROS and insufficient ability of the endogenous antioxidant system to clear ROS, which leads to an increase in hypoxia-inducible factor-1 (HIF-1) levels and further promotes the development of myopia (51, 73). N-omega-nitro-L-arginine methyl ester (L-NAME), an inhibitor of NOS, was injected intravitreally to prevent the development of myopia in both form-deprivation myopia (FDM) and LIM animal models (74, 75). In addition to its anti-inflammatory properties, LF can reverse high levels of MMP-2 activity in the LIM model, thereby inhibiting myopia (63). Additionally, research has indicated that the MMP-9 gene locus may contribute to myopia (76). Combined with our results, the role of macrophages in oxidative stress response may be a key factor affecting the development of myopia.

Currently, studies have shown that some natural compounds or supplements that modulate M1 to M2 macrophages have therapeutic effects on specific diseases. Lupeol, as a multi-target agent, ameliorates experimental inflammatory bowel disease by converting M1 to M2 macrophages (77). Resveratrol through its anti-oxidant capacity and anti-inflammatory properties inhibits platelet aggregation, thereby preventing atherosclerosis (78). Lactobacillus murinus alleviate intestinal ischemia/reperfusion damage by encouraging M2 macrophages to secrete interleukin-10 (79). In addition to all these, there are some compounds that treat ophthalmology-related diseases by their properties of promoting the transformation of M1 into M2 macrophages. Quercetin may have an impact on the treatment of ophthalmic diseases such as conjunctivitis, dry eye, and retinopathy due to its anti-inflammatory and antioxidant properties (80). Forskolin is a medication that can decrease intraocular pressure and be used as an anti-glaucoma agent (81). According to the properties of various natural compounds, which are involved in regulating the phenotype transformation of macrophages from M1 to M2, we targeted the agent named berberine, made it into eye drops, and applied it in myopic-induced mice, we found that it has the ability to inhibit the development of myopia. It indicates a possibility that control of macrophage phenotype may be a new therapy applicable to myopia treatment.

In summary, our results have important implications for understanding the relationship between macrophages and the progression of myopia. In addition to the notion that the two modes of macrophage polarization can have specific effects on choroidal structure as well as the development of myopia, our findings show that the effect of macrophages on myopia development may be related to inflammation, vascular maintenance, and oxidative stress response, and these possible co-existing mechanisms may play a key role in the development of myopia. The above views provide new ideas for regulating the macrophage response to interventions in myopia and provide a new method for drug screening for myopia treatment.

## Materials and Methods

### Mice

The Ethics Committee on Animal Research of the Keio University School of Medicine approved all procedures, which followed the Association for Research in Vision and Ophthalmology Statement for the Use of Animals in Ophthalmic and Vision Research, Keio University’s Institutional Guidelines on Animal Experimentation, and the Animal Research: Reporting of In Vivo Experiments (ARRIVE) guidelines. In addition, this study implemented the principle of random assignment. CLEA Japan, Inc. provided the wild-type male C57BL/6J mice. Those mice were kept in standard transparent mouse cages (29×18×13 cm) with four or five per cage in pathogen-free environments maintained at 23 ± 3 °C, under approximately 50 lux of background fluorescent lamp light (color temperature: 5000 K), with a 12-hour diurnal period and constant access to normal chow (MF, Oriental Yeast Co., Ltd, Tokyo, Japan) and tap water. All the animals were randomly allocated.

### LIM mouse model

A murine LIM model was created as previously described (45). Briefly, mice were placed under general anesthesia using a mixture of midazolam (Sandoz K.K., Tokyo, Japan), medetomidine (Domitor®, Orion Corporation, Espoo, Finland), and butorphanol tartrate (MMB) (Meiji Seika Pharma Co., Ltd., Tokyo, Japan). Eyeglass frames that fit the mouse’s head’s contours were designed and printed using a three-dimensional printer. For myopia induction, a negative 30D lens was constructed using polymethyl methacrylate (PMMA). The left and right eyes of the spectacles were adjusted to fit the shape of the mouse skull frame and screwed onto a stick, which was a joint enabling the left and right frame positions to be altered or removed for cleaning. Subsequently, the stick was bonded to the mouse skull using a self-curing dental adhesive system. Induction began once the mouse had fully recovered from anesthesia, and the lenses were removed for cleaning at least twice a week.

### The measurement of refraction, axial length, and choroidal thickness

Refraction was measured using an infrared photorefractor (Steinbeis Transfer Center, Stuttgart, Baden-Württemberg, Germany) as previously reported (45). After measuring refraction, axial length, and choroidal thickness were measured using an SD-OCT system (Envisu R4310, Leica Microsystems, Wetzlar, Germany). For each mouse, measurements of refraction, axial length, and choroidal thickness were performed before the start of LIM (0W) and at the end of LIM (3W). Before the measurements, the mouse’s eyes were treated with mydriasis eye drops containing 0.5% tropicamide and 0.5% phenylephrine (Santen Pharmaceutical Co., Ltd., Osaka, Japan) to ensure mydriasis and cycloplegia. After pupil dilation, the mice were subjected to general anesthesia using MMB. It is important to prevent the occurrence of corneal injury during measurement. Axial length was measured as the vertical distance from the anterior corneal surface to the retinal pigment epithelium layer near the optic nerve (45). Choroidal thickness was determined by quantifying the circular area of the disc on the posterior surface of the choroid using ImageJ software (82).

### The measurement of choroidal blood perfusion

Choroidal blood perfusion was measured at the beginning (3-week-old) and end (6-week-old) stages of myopia induction using an SS-OCT/OCTA device (XEPHILIO OCT-S1, CANON Medical Systems, Tokyo, Japan). The measurement method has been described in a previous report (83). Briefly, as with refraction measurements, the mice were dilated under general anesthesia before measurement. Choroidal blood perfusion was measured using en face angiography to identify the optic nerve as the central area. The choroidal blood perfusion signal was obtained from B-scan images at the corresponding location. In the B-scan images, the area covered by red noise points was the area without blood perfusion. ImageJ quantitative analysis was used to calculate the non-blood perfusion area in the choroid and evaluate the percentage of the area with blood perfusion in the choroid.

### Flow cytometry

The mice were administered an overdose of MMB to induce profound anesthesia before being euthanized by cervical dislocation. The eyes were immediately enucleated and the anterior segment, vitreous, and retina were discarded. Choroid tissues were carefully scraped off the sclera-choroid complex, collected in the tube (10 choroid tissues/tube), and then processed in digestion buffer (0.75 mg/ml of collagenase A, from FUJIFILM, Tokyo, Japan) in complete Dulbecco’s Modified Eagle Medium (DMEM) solution for 45 min at 37 °C. The digested choroid mixture was transferred onto the cell strainers and any undigested choroid pieces were mashed using the end of a syringe plunger while adding 5 ml cold complete DMEM to allow the pass-through of cells into new 50 ml tubes below the strainer. The aggregated choroidal cells were dissociated into single-cell suspensions via differential centrifugation. The cells were washed by adding 700 ul of flow cytometry staining (FACS) buffer (0.1 g of bovine serum albumin (BSA) and 80 ul of 0.5 mol/L ethylenediaminetetraacetic acid (EDTA) in 20 ml of PBS (pH 7.4) solution to the cell suspension. The pellet was resuspended with 10 ul Fc block solution (1:10 Fc block) (BD Biosciences, New Jersey, USA) for 10 min, stained with a mixture of fluorochrome-conjugated antibodies (1:200 FITC anti-mouse/human CD11b M1/70, BioLegend, California, USA), (1:200 APC/Cyanine7 anti-mouse F4/80, BioLegend, California, USA), (1:200 CD206, AbD Serotec, Kidlington, near Oxford, UK) for at least 45 min, and finally covered with 0.1 mg/ml Hoechst (DOJINDO, Amsterdam, Netherlands) for 10 min. All incubations were performed on ice and light was avoided. Data were acquired on a CytoFLEX S flow cytometer using CytExpert software (Beckman Coulter Life Sciences, Inc., Indiana, United States), and offline data analysis was performed using CyExpert software.

### Clodronate liposome treatment

Clodronate liposome (Liposoma BV, Amsterdam, The Netherlands) was intraperitoneally injected at a dose of 0.10 ml/10 g (once every two days) to C57BL/6J mice from P21 to P28 or from P56 to P63. The control group was injected with an equal amount of PBS liposomes, as described above. Refraction, axial length, and choroidal thickness were measured before and after 4 four injections. The mice were then euthanized and choroid samples were collected.

### LPS treatment

Lipopolysaccharides from *Escherichia coli* O111:B4 (Merck, Tokyo, Japan) were dissolved in PBS and injected intraperitoneally to induce systemic M1 macrophage polarization. The concentration of LPS was 10 mg/ml, and the injection dose was increased proportionally to the body weight of the mice, with 10 μg of LPS given per gram of mice. Since LPS injections caused a large reduction in mouse weight, we monitored and documented the mouse body weight before each injection. In addition, a corresponding volume of PBS was injected intraperitoneally as a vehicle control.

### EPA administration

EPA-supplemented chow containing 5% of EPA (EPADEL®, Mochida, Pharmaceutical Co. Ltd., Tokyo, Japan) and normal chow were prepared as previously reported (43). Briefly, EPA ethyl ester was blended with powdered ordinary chow, and tap water was added. The mixture was shaped into tiny cylinders and dried for consumption. Feeding with EPA-mixed chow began and ended with myopia induction.

### IL-4/IL-13 treatment

To induce systemic M2 macrophage polarization, recombinant murine IL-4 (214-14) and IL-13 (210-13) (PeproTech, New Jersey, United States) were dissolved in PBS at the same concentration of 0.1 µg/100 µl, and injected intraperitoneally into the mice once every two days. The control group was intraperitoneally administered the same volume of PBS. The injections were administered during the induction of myopia. To ensure that the mice grew normally, their body weights were measured and recorded prior to each injection.

### Berberine chloride eye drops

Berberine chloride hydrate (TCI, Tokyo, Japan) solution was freshly prepared in DMSO. When determining the concentration of berberine chloride eye drops, we found that berberine chloride hydrate could not be dissolved in water at room temperature. We attempted to use 100% DMSO to dissolve it and found that the solubility of berberine chloride hydrate in 100% DMSO at room temperature was approximately 13 mg/ml. At present, the concentration of berberine chloride eye drops is 0.26 mg/ml. In addition, DMSO, a dissolution vector, has been reported to exert cytotoxic effects at certain concentrations. Therefore, an untreated control group should be included along with the DMSO vehicle control to check for solvent toxicity (84). To maintain the concentration of berberine chloride hydrate at 0.26 mg/ml and reduce the concentration of DMSO as much as possible, the final concentration of berberine chloride eye drops was 0.26 mg/ml, dissolved in 2% DMSO. Moreover, a parallel control group of 2% DMSO was also set up in mice to observe its effect on myopia.

### Real-time Quantitative PCR (qPCR)

After the mice were euthanized, the eyeballs were enucleated immediately to collect retinal, choroid, and scleral tissues, frozen in liquid nitrogen, and stored at -80 °C for later use. Mouse total RNA was extracted from the retina, choroid, and scleral tissues using TRIzol (TRI reagent) (MOR, Miami, FL, USA #TR118). Small RNAs were isolated using RWT (QIAGEN, Hilden, Germany, #1067933) and RPE buffers (QIAGEN, Hilden, Germany, #1018013). The RNA samples were dissolved in RNase-free water (TAKARA HOLDINGS Inc., Kyoto, Japan, 9012) and measured with a spectrophotometer (NanoDrop; Thermo Fisher Scientific, Waltham, MA, USA). According to the manufacturer’s instructions, the extracted RNA was converted to cDNA after RNA denaturation, DNase reaction, and reverse transcription. mRNA gene expression was determined using SYBR Green RT-PCR on the cDNA template and PCR was performed using StepOnePlus Real-Time PCR System (Applied Biosystems, Waltham, Massachusetts, USA). The mRNA expression of each gene was evaluated using the comparative Ct (ΔΔCT) method and normalized to glyceraldehyde-3-phosphate dehydrogenase (GAPDH) mRNA expression, which was used as the reference gene. The following were the qPCR primer sequences:

mouse *Il6* forward: CTACCCCAATTTCCAATGCT

mouse *Il6* reverse: ACCACAGTGAGGAATGTCCA

mouse *Tnfa* forward: CTGTAGCCCACGTCGTAGC

mouse *Tnfa* reverse: TTGAGATCCATGCCGTTG

mouse Mrc1 forward: TCGAGACTGCTGCTGAGTCCA

mouse Mrc1 reverse: AGACAGGATTGTCGTTCAACCAAAG

mouse *Nox2* forward: ACTCCTTGGAGCACTGG

mouse *Nox2* reverse: GTTCCTGTCCAGTTGTCTTCG

mouse *Nox4* forward: TGAACTACAGTGAAGATTTCCTTGAAC

mouse *Nox4* reverse: GACACCCGTCAGACCAGGAA

mouse *Mmp2* forward: CAAGTTCCCCGGCGAT

mouse *Mmp2* reverse: TTCTGGTCAAGGTCAC

mouse *Mmp9* forward: GGACCCGAAGCGGACA

mouse *Mmp9* reverse: CGTCGTCGAAATGGGC

mouse *GAPDH* forward: AGGAGCGAGACCCCACTAAC

mouse *GAPDH* reverse: GATGACCCTTTTGGCTCCAC

### 8-OHdG ELISA

For assessing oxidative stress, the level of 8-Oxo-2’-deoxyguanosine (8-OHdG) in the choroid tissues was measured using commercial Enzyme-Linked Immunosorbent Assay (ELISA) kits in accordance with the manufacturer’s instructions (Uscn Life Science; Wuhan, China). Because this assay uses a competitive inhibition enzyme immunoassay approach, the concentration of 8-OHdG in the sample and the assay signal strength were inversely correlated. The concentration of 8-OHdG in each sample was calculated using standard curves generated from standard proteins, which were calculated and analyzed using Boster’s ELISA Online Calculator.

### Statistical analyses

Our study used an independent *t*-test and one-way analysis of variance (ANOVA) for statistical significance analyses of all data (Microsoft Excel 2003, USA). All results are presented as mean standard deviation (SD), and results with *P* values < 0.05 were considered statistically significant.

### Study Approval

The Institutional Animal Care and Use Committee at Keio University authorized all operations (approval number: 16017). All methods and experimental protocols followed the National Institutes of Health (NIH) guidelines for working with laboratory animals and the ARVO Animal Statement for the Use of Animals in Ophthalmic and Vision Research.

## Data Availability

All sources of data generated or analyzed during this study are included in this published article and its supplementary information files.

## Author Contributions

Conceptualization, J.H., S.I., T.K., and K.T.; Methodology, J.H., and S.I.; Formal Analysis, J.H.; Investigation, J.H.; Data Curation, J.H.; Project administration, J.H., S.I. and T. K.; Writing – Original Draft Preparation, J.H.; Writing – Review & Editing, S.I., K.M., J.H., H.T., K.N., T.K., and K.T.; Supervision, K.N., T.K., and K.T. All authors made a substantial contribution in the revision of the manuscript.

## Acknowledgments

The authors thank Y. Katada, N. Ban, A. Nakai, N. Serizawa, C. Shoda, M. Ziyan, C. Junhan, and A. Kawabata (Graduate School of Medicine, Keio University, Tokyo, Japan) for their technical guidance and administrative support.

---

**Supplementary Figure 1:**
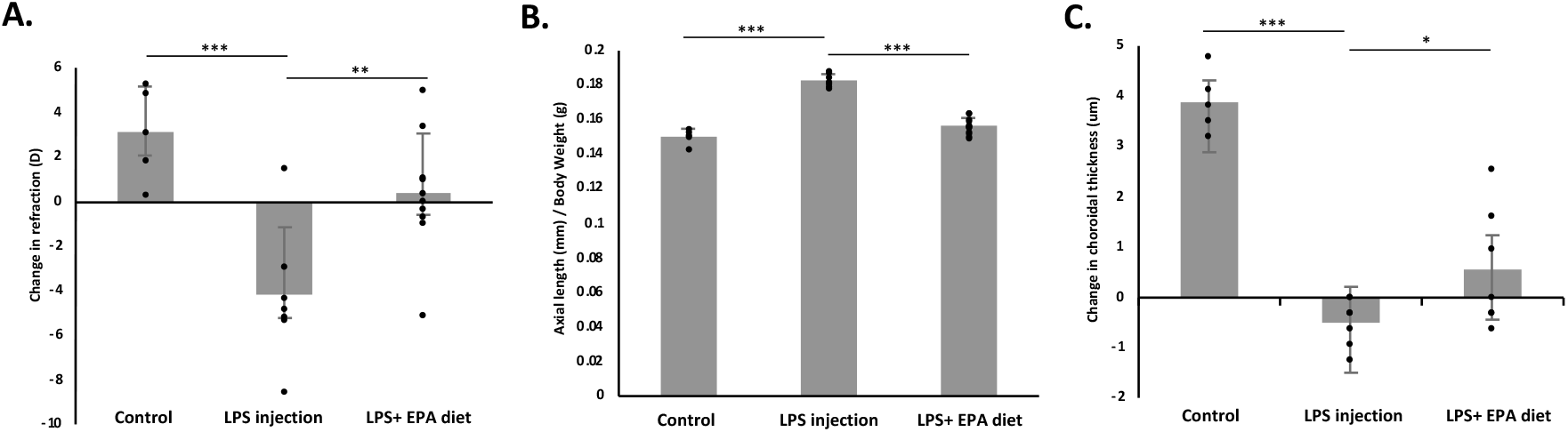
EPA administration inhibited the development of myopia caused by LPS injection. Three-weeks-old wild-type C57BL/6JJc1 mice were divided into three groups: control group, LPS injection group, and LPS administered with EPA group (n=4). Changes in refraction, axial length, and choroidal thickness were measured after 3 weeks of feeding. **(A)** Compared to the large changes in refraction caused by LPS injection, the refractive shift was smaller after EPA administration (*P* < 0.01). **(B)** Compared to the increase in the ratio of axial length to body weight caused by LPS injection, EPA feeding reduced the ratio of axial length to body weight (*P* < 0.001). **(C)** Thinning of choroid occurred in the LPS injection group, whereas EPA feeding significantly improved choroidal thickness (*P* < 0.05). *P* values indicate comparisons relative to the control using one-way ANOVA. \**P* < 0.05. \*\**P* < 0.01. \*\*\**P* < 0.001.

**Supplementary Figure 2:**
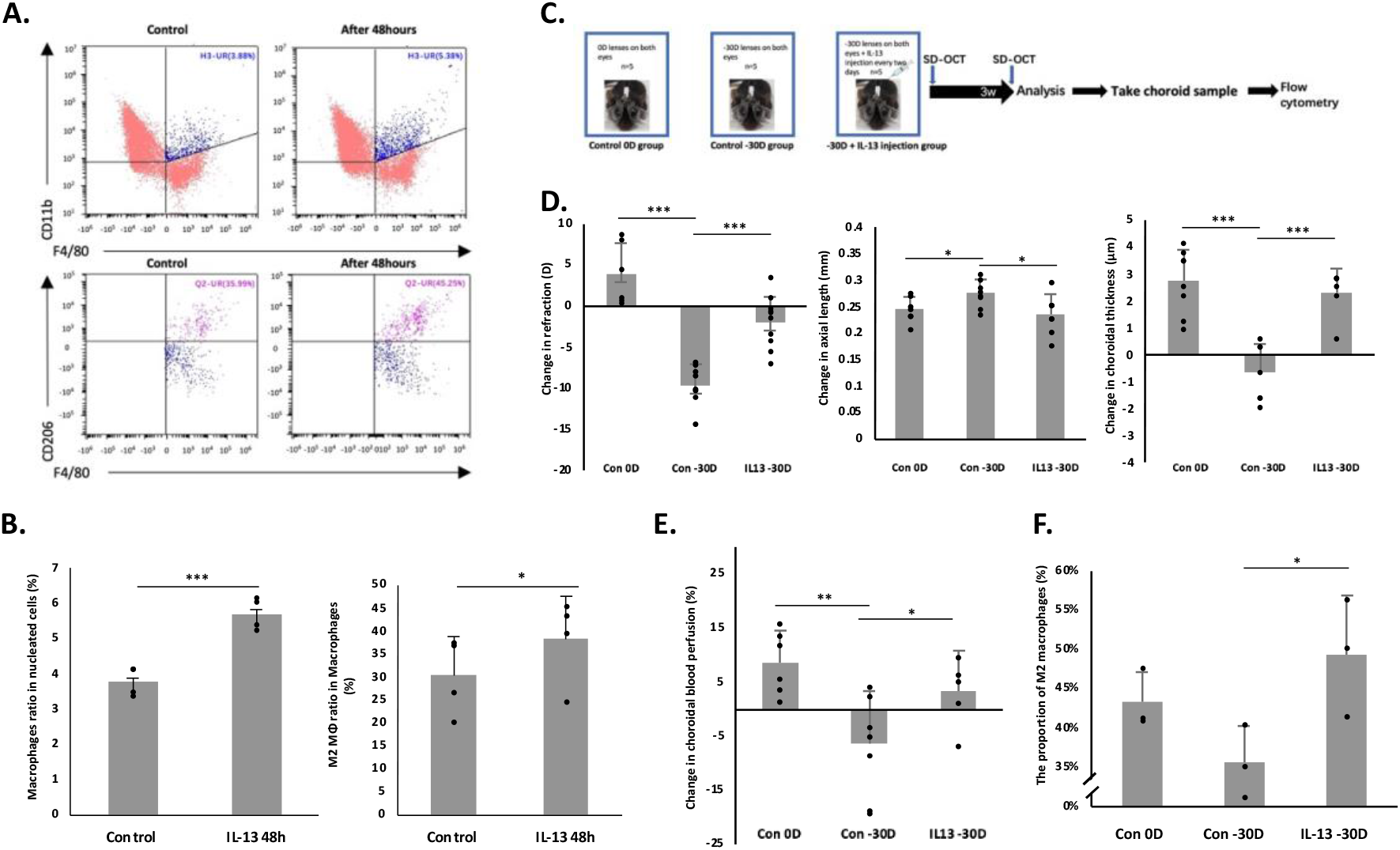
IL-13 injection suppressed myopia progression and increased choroidal blood perfusion in a murine model of lens-induced myopia. **(A)** Choroid samples were collected after 48 h of IL-13 injection and PBS injection for flow cytometry analysis, which was designed to evaluate the optimal time for the polarization of M2 macrophages. **(B)** Flow cytometry analysis demonstrated that both macrophage ratio and M2 macrophage ratio were significantly increased 48 h after IL-13 injection, compared to the PBS injection group. **(C)** Three-week-old mice were divided into three groups: 0D lenses on both eyes (control 0D group), binocular myopia induction (control -30D group), and binocular myopia induction and IL-13 injection (IL-13 -30D group). IL-13 injection frequency was once every two days. Refraction, axial length, and choroidal thickness were measured at the initial (3-week-old) and end (6-week-old) stages of myopic induction with an infrared photorefractor and SD-OCT system. **(D)** The control -30D group showed a significantly larger refractive change (*P* < 0.001), greater axial length elongation (*P* < 0.05), and thinner choroidal (*P* < 0.001) compared to the control 0D group. The IL-13 -30D group indicated a significantly smaller refractive change (*P* < 0.001), lesser axial length change (*P* < 0.05), and positive change in choroidal thickness, with a statistical significance (*P* < 0.001). Bars represent mean +/− standard deviations. **(E)** Choroidal blood perfusion was measured using OCTA. Compared to the control 0D group, choroidal blood perfusion was significantly decreased in the control -30D group. Correspondingly, compared to the control -30D group, choroidal blood perfusion was significantly improved in the IL-13 -30D group. **(F)** The proportion of M2 macrophages in the choroid showed a significant increase in the IL-13 -30D group, compared to the control -30D group. *P* values indicate comparisons relative to control with one-way ANOVA or *t*-test. \**P* < 0.05. \*\**P* < 0.01. \*\*\**P* < 0.001.

**Supplementary Figure 3:**
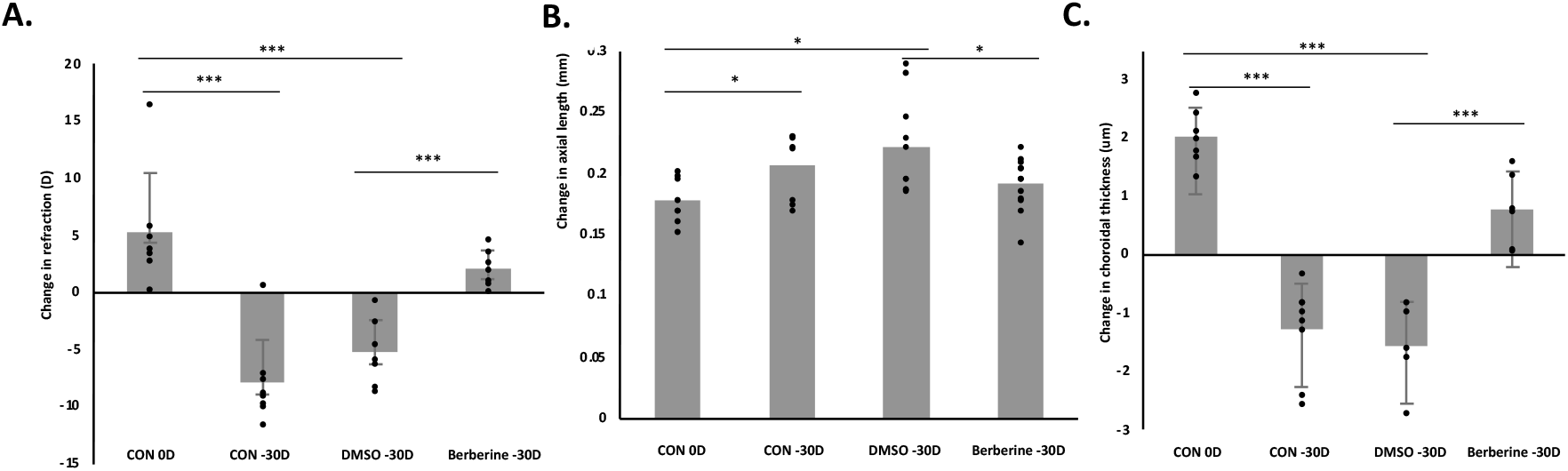
Continuous berberine eye drops can inhibit the progression of myopia in mice. Three-weeks-old wild-type C57BL/6JJc1 mice were separated into control 0D group, control -30D group, 2% DMSO -30D group, and berberine -30D group (n=4). The frequency of eye drops for 2% DMSO and 0.26 mg/ml berberine was once a day. After 3 weeks of eye drops, the changes in refraction, axial length, and choroidal thickness were measured using SD-OCT. **(A)** Compared to the control 0D group, a refractive shift was observed in the control -30D group and 2% DMSO group (*P* < 0.001). Smaller refractive changes were observed in the berberine -30D group compared to the 2% DMSO group (*P* < 0.001). **(B)** Compared to the growth of axial length in the control -30D group and 2% DMSO group (*P* < 0.05), the berberine -30D group showed a smaller axial length elongation (*P* < 0.05). **(C)** In the control -30D group and 2% DMSO -30D group, the choroidal thickness was reduced compared to the control 0D group (*P* < 0.001). Contrarily, choroid thickness was improved in the berberine -30D group compared to the 2% DMSO -30D group (*P* < 0.001). *P* values indicate comparisons relative to the control using one-way ANOVA. \**P* < 0.05. \*\**P* < 0.01. \*\*\**P* < 0.001.

