## Supplemental figures 1~3 for "The orientation of choroidal macrophage polarization significantly influences the development of myopia in murine models": Supplemental Files (BioRxiv).docx

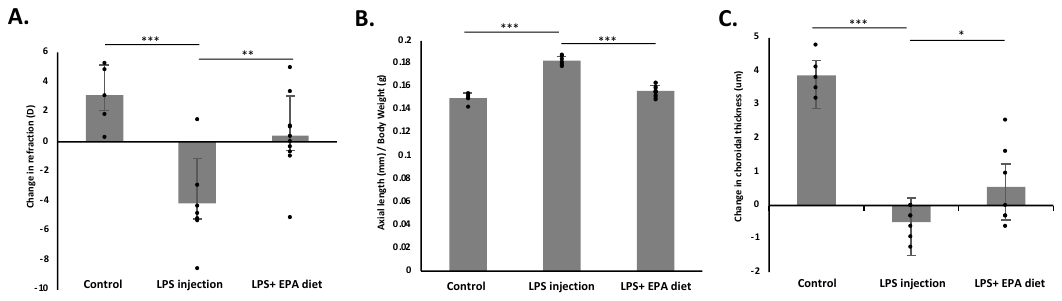

**Supplementary Figure 1: EPA administration inhibited the development of myopia caused by LPS injection.**

Three-weeks-old wild-type C57BL/6JJc1 mice were divided into three groups: control group, LPS injection group, and LPS administered with EPA group (n=4). Changes in refraction, axial length, and choroidal thickness were measured after 3 weeks of feeding. **(A)** Compared to the large changes in refraction caused by LPS injection, the refractive shift was smaller after EPA administration (*P* < 0.01). **(B)** Compared to the increase in the ratio of axial length to body weight caused by LPS injection, EPA feeding reduced the ratio of axial length to body weight (*P* < 0.001). **(C)** Thinning of choroid occurred in the LPS injection group, whereas EPA feeding significantly improved choroidal thickness (*P* < 0.05). *P* values indicate comparisons relative to the control using one-way ANOVA. **P* < 0.05. ***P* < 0.01. ****P* < 0.001.

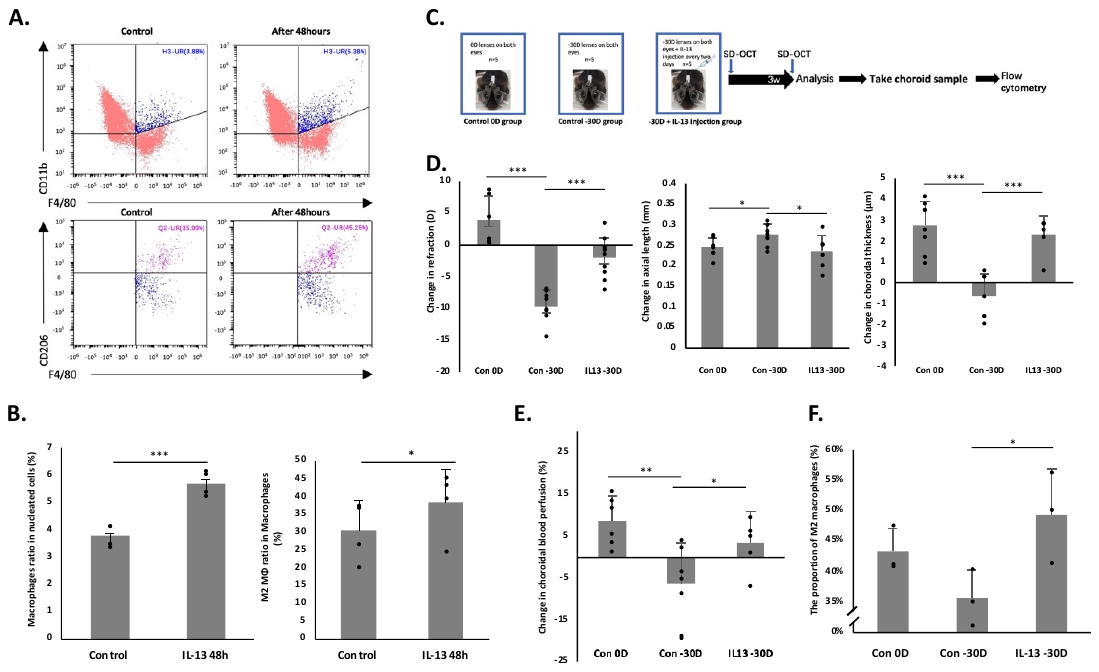

**Supplementary Figure 2: IL-13 injection suppressed myopia progression and increased choroidal blood perfusion in a murine model of lens-induced myopia.**

**(A)** Choroid samples were collected after 48 h of IL-13 injection and PBS injection for flow cytometry analysis, which was designed to evaluate the optimal time for the polarization of M2 macrophages. **(B)** Flow cytometry analysis demonstrated that both macrophage ratio and M2 macrophage ratio were significantly increased 48 h after IL-13 injection, compared to the PBS injection group. **(C)** Three-week-old mice were divided into three groups: 0D lenses on both eyes (control 0D group), binocular myopia induction (control -30D group), and binocular myopia induction and IL-13 injection (IL-13 -30D group). IL-13 injection frequency was once every two days. Refraction, axial length, and choroidal thickness were measured at the initial (3-week-old) and end (6-week-old) stages of myopic induction with an infrared photorefractor and SD-OCT system. **(D)** The control -30D group showed a significantly larger refractive change (*P* < 0.001), greater axial length elongation (*P* < 0.05), and thinner choroidal (*P* < 0.001) compared to the control 0D group. The IL-13 -30D group indicated a significantly smaller refractive change (*P* < 0.001), lesser axial length change (*P* < 0.05), and positive change in choroidal thickness, with a statistical significance (*P* < 0.001). Bars represent mean +/− standard deviations. **(E)** Choroidal blood perfusion was measured using OCTA. Compared to the control 0D group, choroidal blood perfusion was significantly decreased in the control -30D group. Correspondingly, compared to the control -30D group, choroidal blood perfusion was significantly improved in the IL-13 -30D group. **(F)** The proportion of M2 macrophages in the choroid showed a significant increase in the IL-13 -30D group, compared to the control -30D group. *P* values indicate comparisons relative to control with one-way ANOVA or *t*-test. **P* < 0.05. ***P* < 0.01. ****P* < 0.001.

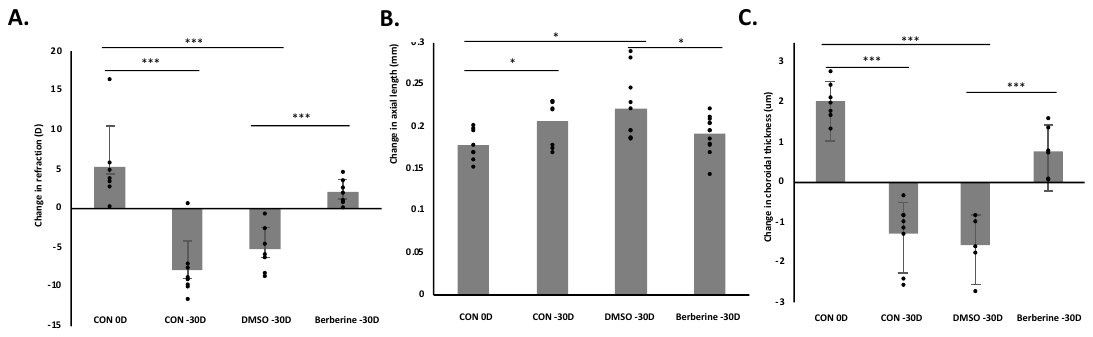

**Supplementary Figure 3: Continuous berberine eye drops can inhibit the progression of myopia in mice.**

Three-weeks-old wild-type C57BL/6JJc1 mice were separated into control 0D group, control -30D group, 2% DMSO -30D group, and berberine -30D group (n=4). The frequency of eye drops for 2% DMSO and 0.26 mg/ml berberine was once a day. After 3 weeks of eye drops, the changes in refraction, axial length, and choroidal thickness were measured using SD-OCT. **(A)** Compared to the control 0D group, a refractive shift was observed in the control -30D group and 2% DMSO group (*P* < 0.001). Smaller refractive changes were observed in the berberine -30D group compared to the 2% DMSO group (*P* < 0.001). **(B)** Compared to the growth of axial length in the control -30D group and 2% DMSO group (*P* < 0.05), the berberine -30D group showed a smaller axial length elongation (*P* < 0.05). **(C)** In the control -30D group and 2% DMSO -30D group, the choroidal thickness was reduced compared to the control 0D group (*P* < 0.001). Contrarily, choroid thickness was improved in the berberine -30D group compared to the 2% DMSO -30D group (*P* < 0.001). *P* values indicate comparisons relative to the control using one-way ANOVA. **P* < 0.05. ***P* < 0.01. ****P* < 0.001.
